# Electrical stimulation of the dorsal motor nucleus of the vagus regulates inflammation without affecting the heart rate

**DOI:** 10.1101/2023.05.17.541191

**Authors:** Aidan Falvey, Santhoshi P. Palandira, Sangeeta S. Chavan, Michael Brines, Kevin J. Tracey, Valentin A. Pavlov

## Abstract

**Background:** The vagus nerve plays an important role in neuroimmune interactions and in the regulation of inflammation. A major source of efferent vagus nerve fibers that contribute to the regulation of inflammation is the brainstem dorsal motor nucleus of the vagus (DMN) as recently shown using optogenetics. In contrast to optogenetics, electrical neuromodulation has broad therapeutic implications, but the anti-inflammatory efficacy of electrical DMN stimulation (eDMNS) was not previously investigated. Here, we examined the effects of eDMNS on heart rate (HR) and cytokine levels in murine endotoxemia as well as the cecal ligation and puncture (CLP) model of sepsis.

**Methods:** Anesthetized male 8–10-week-old C57BL/6 mice on a stereotaxic frame were subjected to eDMNS using a concentric bipolar electrode inserted into the left or right DMN or sham stimulation. eDMNS (50, 250 or 500 μA and 30 Hz, for 1 min) was performed and HR recorded. In endotoxemia experiments, sham or eDMNS utilizing 250 μA or 50 μA was performed for 5 mins and was followed by LPS (0.5 mg/kg) i.p. administration. eDMNS was also applied in mice with cervical unilateral vagotomy or sham operation. In CLP experiments sham or left eDMNS was performed immediately post CLP. Cytokines and corticosterone were analyzed 90 mins after LPS administration or 24h after CLP. CLP survival was monitored for 14 days.

**Results:** Either left or right eDMNS at 250 μA and 500 μA decreased HR, compared with pre- and post-stimulation. This effect was not observed at 50 μA. Left side eDMNS at 50 μA, compared with sham stimulation, significantly decreased serum and splenic levels of the pro-inflammatory cytokine TNF and increased serum levels of the anti-inflammatory cytokine IL-10 during endotoxemia. The anti-inflammatory effect of eDMNS was abrogated in mice with unilateral vagotomy and were not associated with serum corticosterone alterations. Right side eDMNS suppressed serum TNF levels but had no effects on serum IL-10 and on splenic cytokines. In mice with CLP, left side eDMNS suppressed serum TNF and IL-6, as well as splenic IL-6 and increased splenic IL-10 and significantly improved the survival rate of CLP mice.

**Conclusions:** For the first time we show that a regimen of eDMNS which does not cause bradycardia alleviates LPS-induced inflammation and these effects require an intact vagus nerve and are not associated with corticosteroid alterations. eDMNS also decreases inflammation and improves survival in a model of polymicrobial sepsis. These findings are of interest for further studies exploring bioelectronic anti-inflammatory approaches targeting the brainstem DMN.

## Introduction

Inflammation is a localized and timely resolved host response to infection and injury, which is integral to survival. Dysregulated immune responses and excessive inflammation are implicated in the pathogenesis of a broad spectrum of diseases. Discoveries during the last 20 years have revealed the key role of the vagus nerve and the vagus nerve based *inflammatory reflex* in the regulation of cytokine levels and inflammation (1-3). Electrical vagus nerve stimulation of the cervical portion of the vagus nerve has been instrumental in studying the inflammatory reflex. This approach has been shown to significantly suppress serum TNF and other pro-inflammatory cytokine levels in murine endotoxemia and other inflammatory conditions (1, 4). Electrical vagus nerve stimulation has been also successfully explored in treating human inflammatory disorders, including rheumatoid arthritis, and inflammatory bowel disease, and this research has been instrumental for establishing the field of Bioelectronic Medicine (4, 5).

In contrast to this progress, our understanding of the brain mechanisms that govern the inflammatory reflex and vagus nerve anti-inflammatory signalling cholinergic remains limited (6-8). During inflammation, increased levels of peripheral pro-inflammatory cytokine levels trigger activation of neural signaling along afferent (sensory) vagus neurons, which terminate in the brainstem nucleus tractus solitarius (2, 9). This information is communicated to the adjacent dorsal motor nucleus of the vagus (DMN), which is a major source of efferent (motor) vagus cholinergic neurons with anti-inflammatory output as we recently identified using optogenetic stimulation (10). While the future clinical translation of optogenetics, including optogenetic DMN stimulation remains challenging because of the need for expressing light sensitive opsin molecules in the target area, electrical neuronal stimulation has been widely used in brain modulation therapeutic approaches (11). However, whether electrical stimulation can be applied to activate DMN efferent anti-inflammatory signalling remained unknown. The DMN is a major source of efferent preganglionic vagus nerve fibers controlling visceral functions, including cardiac function (12-16). While the functional role of DMN vagal input to the heart is predominantly associated with modulation of the ventricular excitability and contractility, there is some evidence that activation of this input may also cause a decrease in the heart rate (HR) (15-19). Therefore, in addition to studying the effects of electrical DMN stimulation (eDMNS) on inflammation, it has remained imperative to examine whether such effects can be achieved without affecting the HR and avoiding undesirable bradycardia.

Here, we provide new relevant insights. Using eDMNS, we show a current-dependent decrease in the HR in mice. Importantly, we demonstrate for the first time that eDMNS, which does not affect the HR, none-the-less suppresses serum pro-inflammatory cytokine levels in mice with endotoxemia. We also reveal the broader scope of anti-inflammatory effects of this approach and show its therapeutic efficacy in mice with cecal ligation and puncture - a clinically relevant model of polymicrobial sepsis. These findings indicate new possibilities for selective brain immune regulation and inflammatory control using electrical/bioelectronic neuromodulation targeting the DMN and other brain regions.

## Material and methods

### Animals

The Institutional Animal Care and Use Committee and the Institutional Biosafety Committee of the Feinstein Institute for Medical Research, Northwell Health, Manhasset, NY approved all procedures described, in accordance with NIH guidelines. Male, C57BL/6 mice were purchased from Jackson Laboratories aged 7-8 weeks old. All experiments were conducted on mice aged 8-10 weeks old. All mice were given access to food and water *ad libitum* and maintained on a 12h light and dark schedule at 25°C.

### DMN localization and stimulation

Male, C57BL/6 mice were used in eDMNS or sham stimulation experiments. Animals were anesthetized through inhalation of isoflurane gas mixed with air at 2.5%; once anesthetized, mice were transferred to a stereotaxic frame equipped with a nozzle for continuous isoflurane administration at 1–1.5%. A midline skin incision was performed and the underlying muscles retracted exposing the dura matter. The head was tilted forward to ensure adequate viewing of the fourth ventricle through the dura matter. A slit was made in the dura matter with a 23G needle, and the cerebrospinal fluid was cleared and the obex point exposed. A concentric bipolar electrode was fixed to the stereotaxic frame and then placed into the obex point. The electrode was stereotaxically guided to the coordinates of the left DMN (0.25 mm lateral to the obex and 0.48 mm deep to the brainstem surface). In specified instances the right DMN was localized at the same coordinates; however, the electrode was moved in the opposite lateral direction from the obex point. Stimulation was performed, or not (in sham mice), using a multi-channel systems STG4000 series stimulator at 50, 250, or 500 μA, 30 Hz, 260 μsec for five minutes (in experiments with mice with endotoxemia and CLP sepsis). Subsequently, the electrode was withdrawn, and the animal was sutured, removed from the stereotaxic frame and processed further according to the experimental design.

### Heart rate recording

Heart rate was measured prior, during and after eDMNS using the Harvard apparatus - ‘small animal physiological monitoring system’ and external needle electrodes. Recording electrodes were placed sub-dermally in the upper limbs of the mice contralaterally, while a grounding electrode was attached to the lower left limb. The stimulating electrode was guided to the left or right DMN as described. Baseline heart rate was obtained for five minutes, and next electrical stimulation was applied with varying amplitudes (50, 250 & 500 μA) for one minute.

### Unilateral vagus nerve section

In select instances, a unilateral left vagus nerve neurotomy (transection) was performed prior to the described eDMNS. Mice were anesthetized with 2.5% isoflurane gas mixed with air and maintained at 1.5%. A cervical midline incision was performed, and the underlying muscle was carefully retracted. The left vagus nerve was identified, and the carotid sheath removed. The left vagus nerve was isolated and transected solely to avoid damage to surrounding tissue and mechanical activation. In sham animals the carotid sheath was removed, and the left vagus nerve identified. Animals were sutured and allowed to recover for one week prior to eDMNS as previously described.

### Endotoxemia

Directly prior to placing the mice in the recovery cage, lipopolysaccaharide (LPS, endotoxin, Sigma-Aldrich) was administered intraperitoneally (0.5 mg/kg) and mice were allowed to recover in a heated cage for 90 minutes prior to euthanization via CO2 asphyxiation and blood through cardiac puncture and the spleen were collected and processed for further analyses.

### Cecal ligation and puncture (CLP)

Cecal ligation and puncture (CLP) was performed in a cohort of C57BL/6, male mice, aged 10-12 weeks. Mice were anesthetized using isoflurane gas mixed with air at 2.5%; once anesthetized mice were maintained at 1.5%. An abdominal midline incision was made, followed by a laparotomy. The cecum was identified and partially removed through the opening, a suture was tied around the length of the cecum and pierced with a 22-gauge needle twice, the cecum was subsequently gently squeezed to protract a measure of stool from the piercings. The cecum was returned to the abdominal cavity and the animal was sutured. Subsequently the animal was flipped and eDMNS or sham stimulation was performed as described above. Animals were recovered in a heated cage and then returned to their home cage for 24 hours until euthanization by CO2 asphyxiation. Blood was withdrawn through cardiac puncture and centrifuged for serum analysis. Additionally, the spleen was removed and processed in tissue protein extraction reagent for cytokine analysis and protein quantification.

### Analyte measurement

Blood was withdrawn and centrifuged for serum analysis. The spleen was removed and processed in tissue protein extraction reagent for cytokine analysis and protein quantification using a standard Bradford protein assay kit (Biorad). Subsequently tissues and serum were analyzed for cytokine levels using standard ELISA kits: TNF (88-7324: Invitrogen); IL-6 (R&D Systems DY406); and IL-10 (R&D Systems DY417 and Luminex). Serum cytokines were expressed as picogram per milliliter of serum collected and splenic cytokines were expressed as picograms per mg splenic protein.

### Statistics

Heart rate data is expressed as pooled traces over time ± standard error of the mean: for quantification, area under the curve was found for each individual trace (pre-, during-, & post-stimulation) and then standardized to the pre-stim values – a paired one-way ANOVA was subsequently performed. All cytokine data is presented as means ± standard error of the mean. All individual points refer to one animal, typically data is pooled from multiple small cohorts - n is expressed for each figure. Data was analyzed for normality using a Shapiro-Wilk normality test. If normal; data was processed for Student’s t-test or ordinary one-way ANOVA or two-way ANOVA using GraphPad Prism 9, all specific statistical tests are listed in the figure legends.

## Results

### Electrical DMN stimulation which suppresses inflammation can be accomplished without affecting heart rate

Endotoxemia, induced by peripheral lipopolysaccharide (LPS, endotoxin) administration provides a standardized model to study innate immune responses and the inflammatory and cardiometabolic effects of systemic cytokine release both in animals and humans (20, 21). Recently, in ChAT-ChR2-eYFP mice we showed that optogenetic stimulation of the left DMN, which contains a substantial cluster of cholinergic neurons projecting within the efferent vagus nerve, suppresses serum TNF levels during endotoxemia (10). As there is some evidence that DMN stimulation using optogenetic stimulation and other approaches may also decrease the HR (14-16, 18), and bradycardia is an undesirable side effect, here we studied whether eDMNS can be applied to selectively alter cytokine levels in endotoxemic mice in a manner that does not affect the HR. We first examined the effects of three different regimens of eDMNS in which 30 Hz and 260 μsec were kept constant and combined with 500 μA, 250 μA, or 50 μA on the HR. Briefly, anesthetised C57BL/6 mice were placed on the stereotaxic frame, and surgical intervention was performed to slowly insert a concentric bipolar electrode in the left DMN using previously established coordinates (10). The HR was recorded for 1 min prior, during, and post-stimulation. As shown in **Figure 1A and B**, eDMNS significantly inhibited the HR using a current of 500 μA, there was a trend towards lower HR at 250 μA, and no significant effect was observed when 50 μA were used. Based on the HR results, in other cohorts of mice, we studied the effect of eDMNS on cytokine responses in mice subjected to endotoxemia (**Figure 1C**). Sham stimulation (electrode inserted, but no electricity) or eDMNS was carried out prior to administering LPS (i.p.) and the animals were euthanized 90 mins later. We first examined whether a regimen of eDMNS that altered (albeit not significantly) the HR - 30 Hz, 260 μsec, 250 μA, for 5 mins would affect serum TNF levels. As shown in **Figure 1D**, eDMNS compared with sham stimulation resulted in a significant decrease in serum TNF. Then, in another cohort of mice we applied sham stimulation or a regimen of eDMNS, which did not cause any alterations in the HR – i.e., 30 Hz, 260 μsec, 50 μA, for 5 mins prior to subjecting mice to endotoxemia. The lack of effect of this regimen of eDMNS on the HR was confirmed for the entire duration (5 min) of stimulation compared with pre- and post-stimulation (**Supplementary Figure 1**). eDMNS, compared with sham stimulation significantly decreased serum TNF levels (**Figure 1E)**. This result was indicative for the possibility to beneficially alter cytokine responses without affecting the HR. Therefore, to broaden the insight, we also examined the effect of eDMNS on the serum levels of the anti-inflammatory IL-10 (22). As shown in **Figure 1E**, serum levels of the anti-inflammatory cytokine IL-10 were significantly increased in the eDMNS group compared with sham stimulated controls. As the spleen is a major target organ of the efferent arm of the inflammatory reflex (23, 24), we also studied the effects of left side eDMNS on splenic cytokine levels. As shown in **Figure 1F**, eDMNS significantly decreased splenic TNF levels and had no significant effect on splenic IL-10 levels. These results indicate that applying eDMNS can cause bradycardia, but an eDMNS regimen that does not alter the HR can significantly reduce inflammation in mice with endotoxemia.

**Figure 1.**
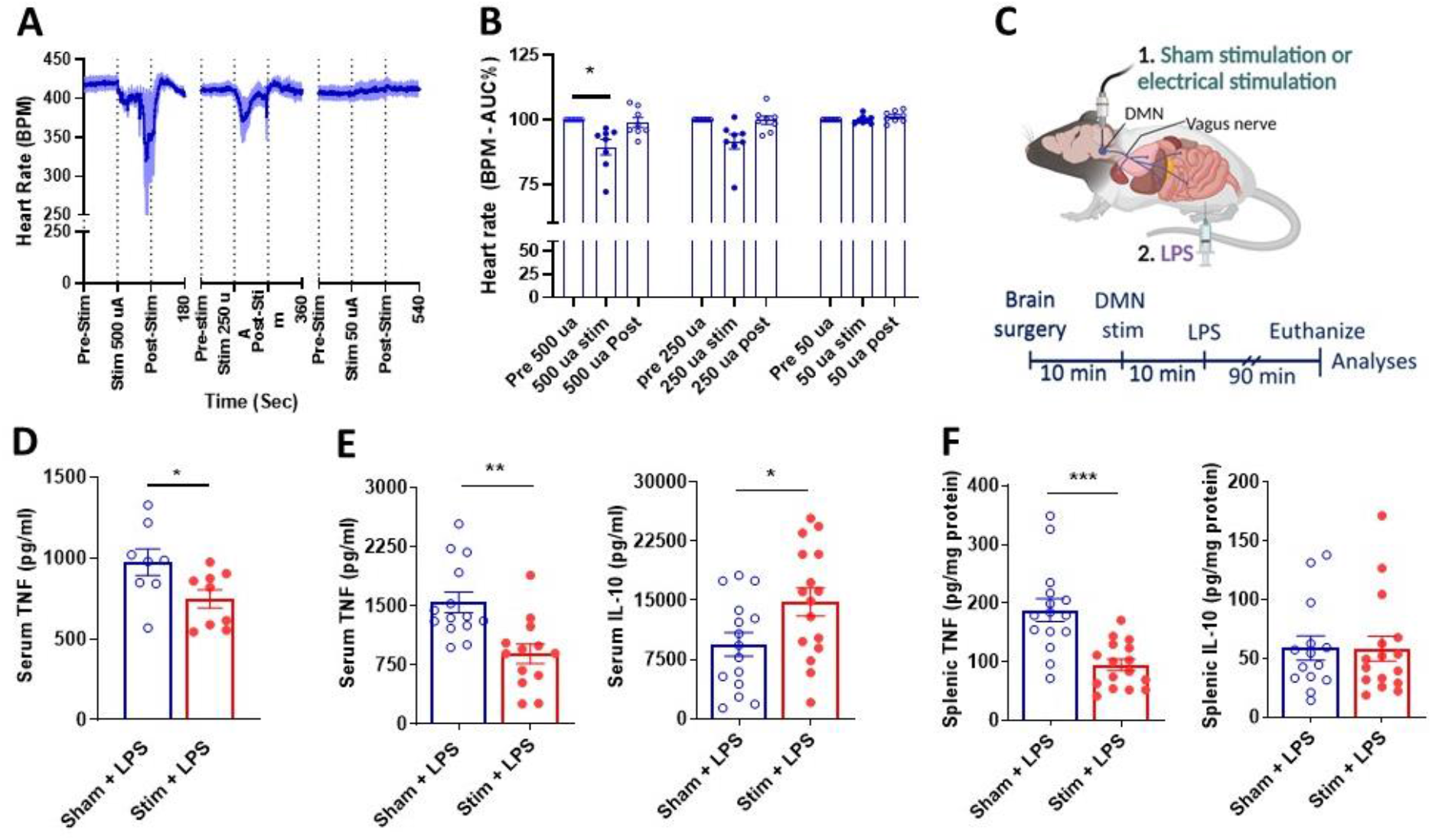
Electrical left side DMN stimulation (eDMNS) regulates the heart rate and alters serum and splenic cytokines in endotoxemic mice. eDMNS causes a current dependent decrease in the heart rate: **(A)** HR recordings; (**B)** Area under the curve (AUC) analysis (*P = 0.031). Data are represented as individual mouse data points with mean ± SEM. One-way ANOVA was used with Dunn’s multiple comparisons. (**C**) Schematic depiction of the experimental design using eDMNS in endotoxemic mice. Anesthetized mice were placed on a stereotactic frame and following a surgical intervention to insert an electrode in the left DMN and sham stimulation or eDMNS was performed for 5 mins and LPS (0.5 mg/kg, i.p.) was administered. Mice were euthanized 90 mins later and blood and spleen were collected and processed for analysis. (**D**) eDMNS (30 Hz, 260 μsec, 250 μA) compared with sham stimulation significantly decreases serum TNF (**P = 0.037; Students *t*-test), (**E**) eDMNS (30 Hz, 260 μsec, 50 μA) significantly decreases serum TNF (**P = 0.001; Students *t*-test) and increases IL-10 levels. (*P = 0.021; Students *t*-test) (**F**) eDMNS (30 Hz, 260 μsec, 50 μA) significantly suppresses splenic TNF (***P = 0.0001; Students *t*-test) but does not significantly alter splenic IL-10. Data are represented as individual mouse data points with mean ± SEM. (See Material and Methods for details).

### Electrical DMN stimulation does not suppress TNF levels in vagotomised mice and does not alter serum corticosterone levels during endotoxemia

We next assessed the role of the vagus nerve in mediating the anti-inflammatory effect of eDMNS. Mice were subjected to surgical transection of the left vagus nerve (unilateral vagotomy) or sham operation one week prior to applying sham stimulation or left side eDMNS in the two groups of mice. As shown in **Figure 2A**, eDMNS compared with sham stimulation significantly suppressed serum TNF levels in sham operated control mice; however, this anti-inflammatory effect was abrogated in mice with ipsilateral vagotomy. Similarly, while eDMNS significantly suppressed splenic TNF levels in sham operated controls, the effect was abolished in mice with vagotomy (**Figure 2B**). Corticosteroids as a result of activation of the hypothalamic pituitary adrenal axis by exogenous neuronal stimuli can be involved in the regulation of inflammation (4, 25). Therefore, to examine whether eDMNS may alter this axis, we also assessed the effects of left side eDMNS on serum corticosterone levels. As shown on **Figure 2C**, there was no significant difference in these levels between sham stimulated mice and mice subjected to eDMNS. These results confirm a critical role of the vagus nerve in mediating the anti-inflammatory effect of eDMNS and suggest that no activation of a neuroendocrine mechanism with altered corticosteroid release is associated with eDMNS in our experimental procedures.

**Figure 2.**
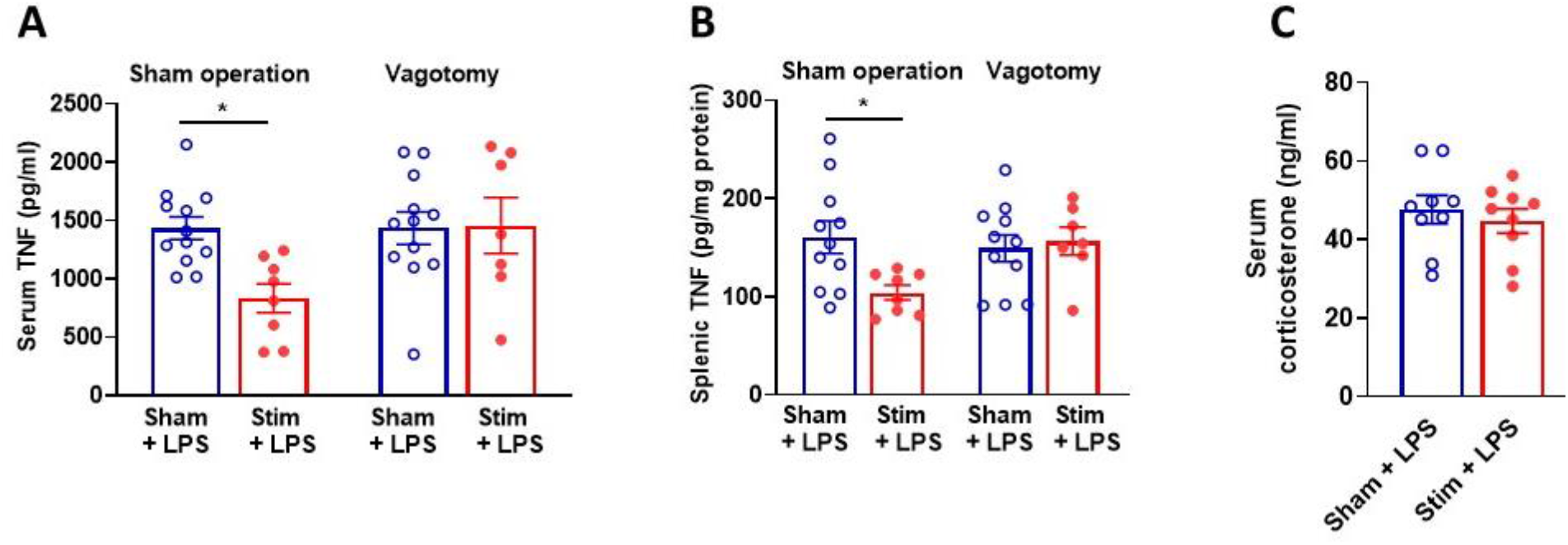
The anti-inflammatory effect of electrical left side DMN stimulation (eDMNS) is abrogated in mice with ipsilateral cervical vagotomy and is not related to corticosteroid level alterations during endotoxemia. eDMNS, compared with sham stimulation significantly suppresses (**A**) serum TNF (*P = 0.012) and (**B**) splenic TNF (*P = 0.012) in mice with sham operation and these effects are abolished in mice with unilateral cervical vagotomy during endotoxemia. (**C**) eDMNS, compared with sham stimulation does not significantly change serum corticosterone levels during endotoxemia. Data are represented as individual mouse data points with mean ± SEM. Two-way ANOVA was used with Sidak’s multiple comparisons (See Material and Methods for details).

### Electrical right side DMN stimulation suppresses serum TNF levels without affecting heart rate

While the left vagus nerve stimulation has been predominantly utilized in treating inflammation, there is also evidence for anti-inflammatory effects of right VNS (4, 26). Therefore, we also examined the possibility to alter cytokine responses via applying right side eDMNS. As shown in **Figure 3A and B**, right side eDMNS resulted in regimen-dependent HR suppression; while 500 μA or 250 μA caused significant HR suppression, 50 μA did not alter the HR. Accordingly, we next examined the effect of right side eDMNS using 30 Hz, 260 μsec, 50 μA, for 5 mins on cytokine responses in endotoxemic mice. Applying this eDMNS resulted in significant suppression in serum TNF levels but did not alter serum IL-10 (**Figure 3C**). Moreover, no significant differences in splenic cytokines between the stimulated and the sham stimulated groups of mice were found (**Figure 3D**). These results indicate that applying right side eDMNS can cause bradycardia, but an eDMNS regimen that does not affect the HR can significantly reduce serum TNF levels without significantly affecting IL-10 and the splenic cytokine levels.

**Figure 3.**
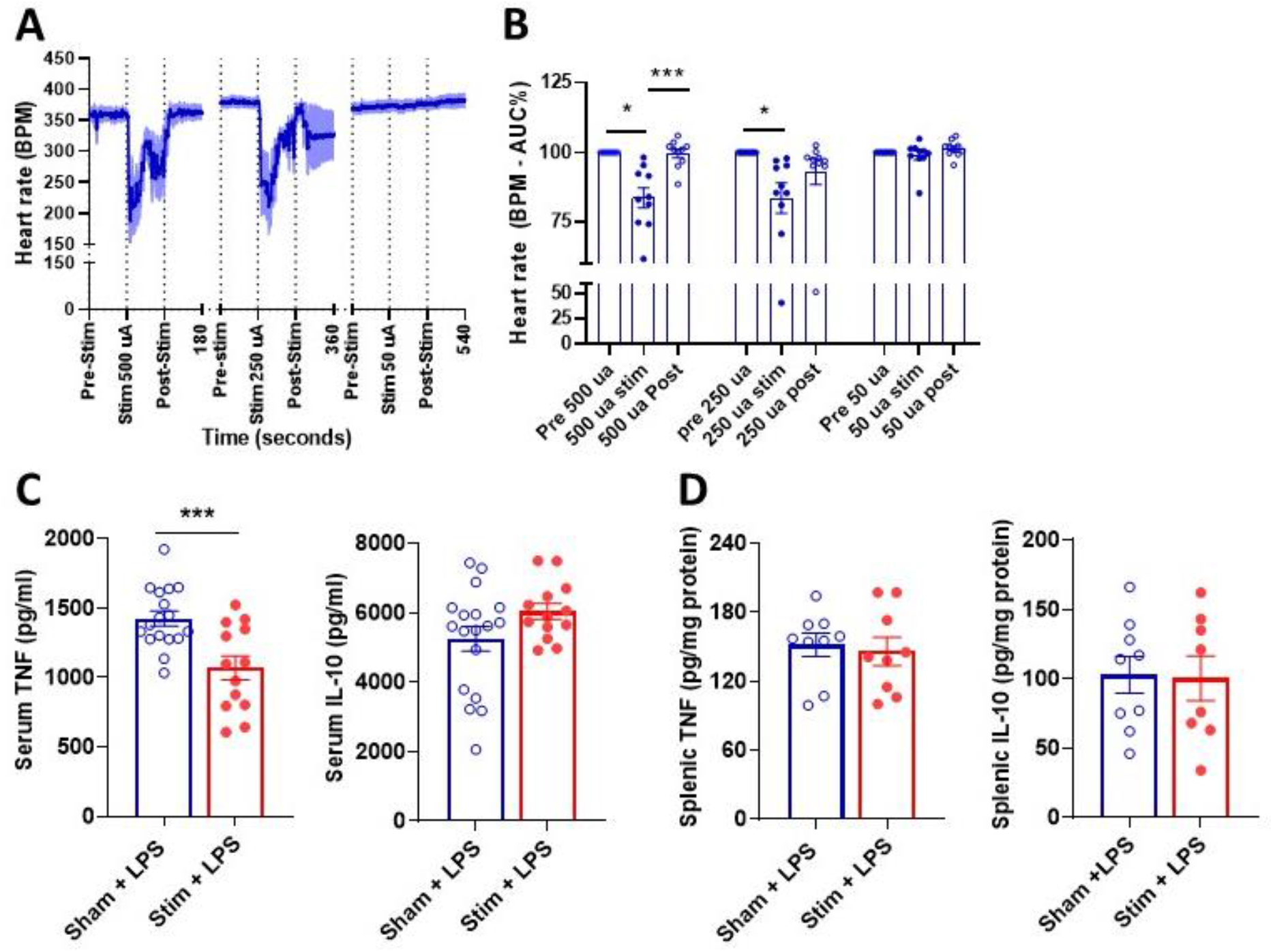
Electrical right side DMN stimulation (eDMNS) regulates the heart rate and alters serum cytokines in endotoxemic mice. eDMNS causes a current dependent decrease in the heart rate: **(A)** HR recordings; (**B)** Area under the curve (AUC) analysis (*P = 0.022 (Pre vs 500 μA Stim); ***P = 0.0008 (500 μA Stim vs Post); *P = 0.039 (Pre vs 250 μA Stim)) Data are represented as individual mouse data points with mean ± SEM. One-way ANOVA was used with Dunn’s multiple comparisons. (**C**) eDMNS (30 Hz, 260 μsec, 50 μA) significantly decreases serum TNF (***P = 0.001; Students *t*-test) and does not alter serum IL-10 levels. (**D**) eDMNS (30 Hz, 260 μsec, 50 μA) does not significantly alter splenic TNF and IL-10. Data are represented as individual mouse data points with mean ± SEM. See Material and Methods for details.

### Electrical DMN stimulation alters serum and splenic cytokine levels and improves survival in mice with CLP-induced polymicrobial sepsis

While endotoxemia is a model of systemic inflammation arising from gram negative sepsis (21), cecal ligation and puncture (CLP) is a clinically relevant and widely used model of polymicrobial sepsis (21). Therefore, to broaden the insight on the therapeutic utility of eDMNS we studied the effects of this approach in mice subjected to CLP and based on our results in the endotoxemia model, we studied the anti-inflammatory efficacy of left eDMNS during CLP. As depicted in **Figure 4A**, we examined the therapeutic effects of eDMNS on cytokine responses, implementing an experimental design in which animals were first subjected to CLP. Briefly, C57BL/6 male mice were anesthetised, the CLP surgery was performed, and the animals were sutured and then placed into a stereotaxic frame for a surgical intervention to apply sham stimulation or eDMNS (50 μA, 30 Hz, 260 μsec, for 5 mins) as described in detail in Material and Methods. Twenty-four hours after the CLP, mice were euthanized, and blood and spleen were collected and processed for cytokine analysis. As shown in **Supplementary Figure 2**, eDMNS (compared with sham stimulation) significantly supressed serum TNF and decreased (albeit not significantly) splenic TNF levels in mice with CLP. While in endotoxemia, TNF is validated indicator of systemic inflammation, the role of TNF in CLP pathogenesis has not been definitively established. However, the pro-inflammatory cytokine IL-6 has proven useful as a marker of sepsis severity in preclinical settings and in humans (27, 28) and a therapeutic target in experimental sepsis (29). Therefore, we also analysed the effect of eDMNS on serum and splenic IL-6 and IL-10 levels in mice with CLP sepsis. As shown on **Figure 4B**, eDMNS (compared with sham stimulation) significantly supressed serum IL-6 levels and did not significantly change serum IL-10 levels. eDMNS (compared with sham stimulation) also significantly suppressed splenic IL-6 levels and increased splenic IL-10 levels (**Figure 4C**). In addition, eDMNS (compared with sham stimulation) significantly improved the 14-day survival rate of mice subjected to CLP. These results demonstrate the differential, cytokine specific anti-inflammatory effects of eDMNS in mice with CLP-induced polymicrobial sepsis and the efficacy of this approach in improving CLP survival.

**Figure 4.**
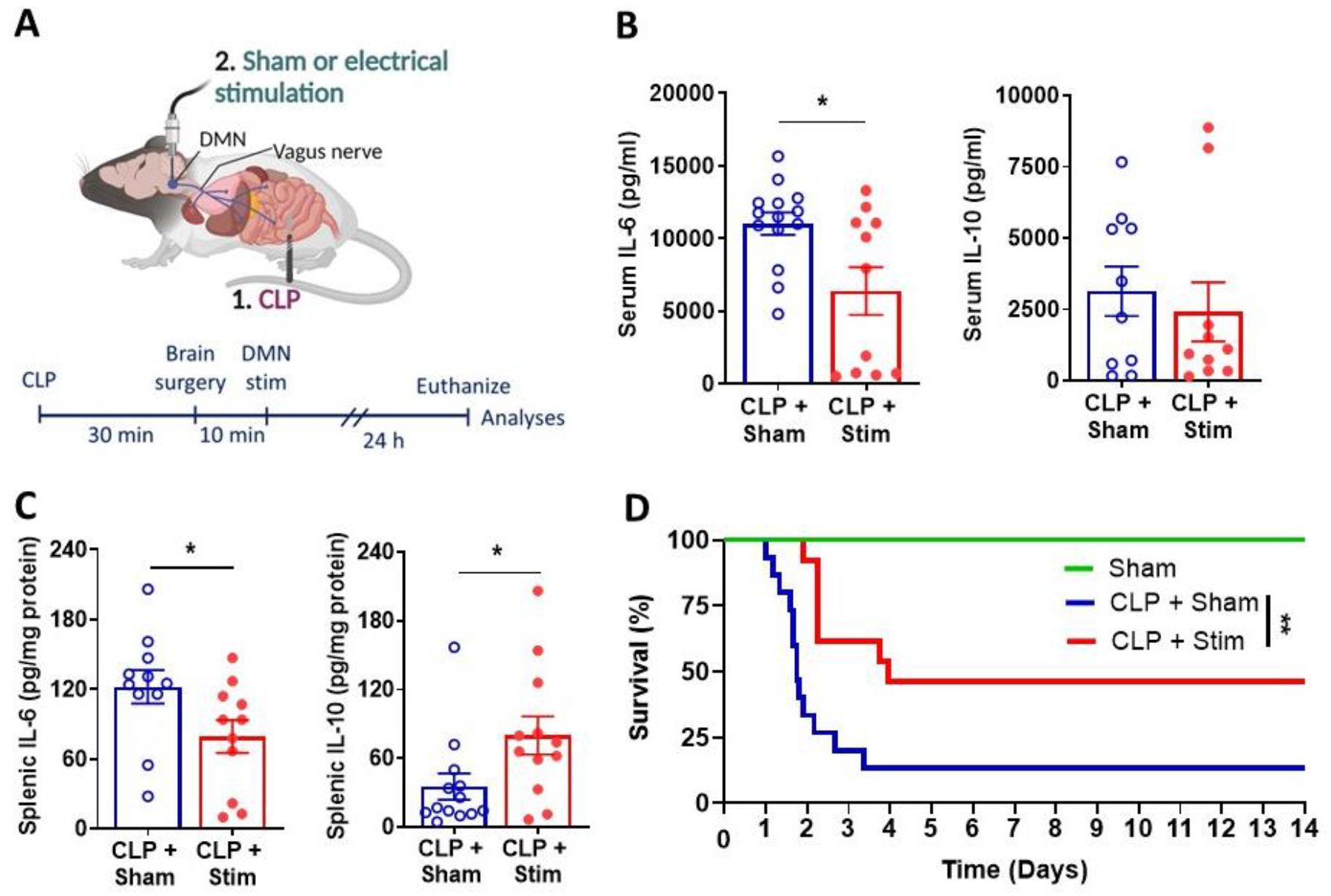
Electrical left side DMN stimulation (eDMNS) suppresses inflammation and improves survival in mice with CLP sepsis. **(A)** Schematic depiction of the experimental approach for eDMNS in mice with CLP sepsis. eDMNS or sham stimulation was performed in anesthetized mice after CLP surgery and cytokines at 24h, or 14-day survival were analyzed. (**B**) eDMNS (30 Hz, 260 μsec, 50 μA) significantly decreases serum IL-6 (*P = 0.0384; Mann-Whitney test) and has no effect on IL-10 levels. (**C**) eDMNS significantly decreases splenic IL-6 (*P = 0.047; Students *t*-test) and increases IL-10 (*P = 0.035; Mann-Whitney test) levels. Data are represented as individual mouse data points with mean ± SEM. Unpaired *t* test was used. (See Material and Methods for details). (**D**) eDMNS compared with sham stimulation (30 Hz, 260 μsec, 50 μA) significantly improves survival (**P = 0.004, Mantel-Cox Test) See Material and Methods for details.

## Discussion

Here, we show for the first time that electrical stimulation can be applied to the brainstem DMN to beneficially alter inflammatory indices in mice with endotoxemia and CLP-induced sepsis without potentially deleterious effects on the HR.

The vagus nerve plays a major role in the regulation of cytokine levels and inflammation (1-3, 30-33). Electrical stimulation of the cervical vagus nerve, which suppresses aberrant inflammation in murine models of endotoxemia and other inflammatory conditions (1, 8, 24, 34-36) has been successfully explored in the clinical treatment of inflammatory and autoimmune diseases (1, 4). The vagus nerve contains afferent (about 80%) and efferent (about 20 %) fibers and cervical VNS stimulates both types, which decreases selectivity and theoretically increases the possibility of side effects. The brainstem DMN is a major source of efferent vagus fibers and provides a locus for targeted stimulation of these neurons in anti-inflammatory exploration. Using optogenetic stimulation we recently revealed the role of DMN cholinergic neurons in suppressing serum TNF levels during murine endotoxemia (10). This study and other reports employing optogenetic stimulation emphasize the importance of studying targeted brain neuromodulation in new strategies for controlling inflammation (6, 37-39). While optogenetic stimulation provides a cell specific means of neuromodulation, the clinical applicability of this approach remains challenging. In contrast, there is a great progress in using electrical stimulation to modulate specific brain areas for therapeutic benefit (40, 41). For instance, deep brain stimulation is a standard of care for Parkinson’s disease, essential tremor and dystonia (40, 41) and this approach targeting various brain regions has been explored in the treatments of epilepsy, Alzheimer’s disease and other disorders (42). However, the possibility of applying electrical stimulation on the DMN to control cytokine responses and inflammation remained unexplored.

Among other organs, DMN efferent vagus nerve fibers innervate the heart and regulate cardiac function (12-16). The physiological role of DMN vagal cardiac innervation is manly linked to modulation of the ventricular excitability and contractility (17-19). However, chronotropic effects as a result of activation of DMN cardiac projections have also been reported (14-16). Therefore, prior to investigating its effects on inflammation we examined the effects of eDMNS on the HR, because desirable anti-inflammatory effects of eDMNS should not be associated with undesirable bradycardia. Altering the stimulation parameters, we show a current-dependent effect of left eDMNS on the HR and the lack of effect when a current of 50 μA is used. Then in mice with endotoxemia we show that this eDMNS significantly suppresses serum TNF and increases the levels of the anti-inflammatory cytokine IL-10 compared with sham stimulation. In addition, eDMNS significantly decreases TNF in the spleen – a major target organ of the efferent arm of the inflammatory reflex, in which the efferent vagus nerve interacts with the splenic nerve in the celiac superior mesenteric ganglion complex (10, 43-46).

Our results obtained in experiments with mice subjected to vagotomy or sham surgery prior to eDMNS clearly indicate that the anti-inflammatory effect of eDMNS is mediated through the vagus nerve, because ipsilateral vagotomy abrogated the effect. However, eDMNS does not change serum corticosteroid levels, which indicates that activation of brain derived neuroendocrine mechanisms such as the hypothalamic pituitary adrenal axis does not play a mediating role.

We also demonstrate the current-dependent effects of right side eDMNS on the HR and the lack of effect at 50 μA. This eDMNS regimen, which does not affect the HR also results in anti-inflammatory effects. However, in contrast to left eDMNS, these effects do not involve alterations in the serum levels of the anti-inflammatory cytokine IL-10, and changes in the splenic cytokines analyzed. These results reveal a differential, side dependent DMN anti-inflammatory regulation using the same parameters of eDMNS. They also indicate that suppressing serum TNF and IL-6 using left eDMNS occurs in parallel with and may be related to decreasing the splenic levels of these cytokines. However, the suppression of serum TNF using right DMNS is not associated with decreases in the splenic levels of this cytokine - an observation that suggests that suppression of cytokine production and release in other organs targeted through right eDMNS may play a role. The increase in the serum levels of IL-10 through left eDMNS and the lack of significant effect of right DMN is a novel observation, which also points to side-dependent DMN differential regulation of this prototypical anti-inflammatory cytokine (22). These observations are in line with previous reports indicating differences in anti-inflammatory regulation applying right cervical VNS in other murine models of inflammatory disorders (26).

Our results demonstrating the effects of eDMNS in settings of experimental sepsis substantially broaden the insight about the therapeutic efficacy of this approach. For the first time, we show that eDMNS alters serum cytokines and improves survival in mice with CLP – a widely used clinically relevant model of polymicrobial sepsis. Left side eDMNS applied after CLP suppresses serum TNF and IL-6 – effects observed also in mice with endotoxemia. However, in contrast to endotoxemia, eDMNS does not significantly affect serum IL-10 levels in CLP mice. Interestingly, while the reduction in serum IL-6 is associated with lower splenic IL-6 following eDMNS, there is also a significant increase in splenic IL-10 levels in the stimulated group, which does not reflect changes in the serum levels of this anti-inflammatory cytokine. Importantly, eDMNS does not merely influence serum and splenic cytokine levels, but also results in significant survival improvement in this lethal CLP model. These effects indicate the therapeutic utility of a single eDMNS performed after CLP in this experimental model of sepsis and rationalize performing future studies utilizing other timeframes and paradigms of stimulation that will provide additional relevant insight.

Sepsis is a complex disease defined as life-threatening organ dysfunction caused by a dysregulated host response to infection (47). Sepsis affects approximately 1.7 million adults in the United States each year and is linked to more than 250 000 deaths, which makes it a leading cause of death in the US hospitals and a major public health problem (48-50). Currently, there are no clinically approved specific treatments for sepsis, which dictates the need for evaluating new therapeutic approaches. The brain is also affected during sepsis and cognitive deterioration, delirium and comma have been documented within the scope of sepsis-associated encephalopathy (51, 52). Accordingly, the brain in sepsis has been mostly evaluated in terms of possibilities to treat these neuropsychiatric manifestations. Our results indicate that the brain also provides a therapeutic target for controlling cytokine responses in sepsis and improving the disease survival.

## Conclusion

Our results reveal that during inflammation, eDMNS parameters which do not cause bradycardia, none-the-less beneficially alters cytokine responses through signaling that requires an intact vagus nerve and does not affect corticosteroid levels. The results also indicate the broad scope of therapeutic anti-inflammatory efficacy of eDMNS in a preclinical scenario of a complex disease such as sepsis. These findings are of interest for enabling future bioelectronic approaches targeting DMN and other brain regions for therapeutic benefit in conditions characterised by immune dysregulation and excessive inflammation.

## Acknowledgements

This work was supported by the National Institutes of Health (NIH), National Institute of General Medical Sciences grants: RO1GM128008 and RO1GM121102 (to VAP) and R35GM118182 (to KJT).

## Author contributions

AF and VAP designed research. AF and SP performed research. AF, SP, and VAP analyzed and interpreted data. AF and VAP wrote the manuscript. MB, SSC, and KJT provided additional comments to finalize the paper.

## Data availability statement

Data is available at the authors’ discretion upon direct request to the corresponding authors.

