## Supplementary Figures 1 and 2 for "Electrical stimulation of the dorsal motor nucleus of the vagus regulates inflammation without affecting the heart rate"

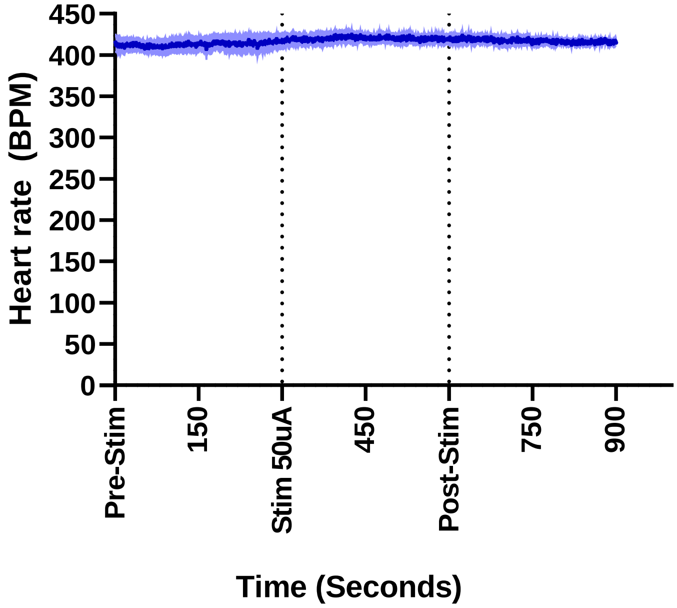


**Supplementary Figure 1. Electrical left side DMN stimulation (eDMNS) (30 Hz, 260 μsec, 50 μA, for 5 mins) does not alter the heart rate** (HR recordings of 4 mice).

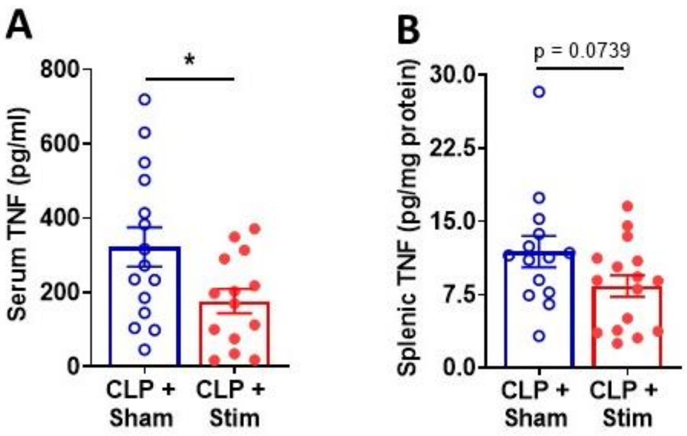


**Supplementary Figure 2. Electrical left side DMN stimulation (eDMNS) (A) suppresses serum TNF levels and (B) does not significantly affect splenic TNF in mice with CLP sepsis.** Data are represented as individual mouse data points with mean ± SEM. *P = 0.0291, unpaired Student *t*- test. See Material and Methods for details.
